# Multiscale modelling of drug–host–pathogen interaction: quantifying drug and immune contributions to treatment response

**DOI:** 10.64898/2026.08.30.744418

**Authors:** Alessandro Ravoni, Enrico Mastrostefano, Davide Moretti, Elia Onofri, Francesca Pelusi, Aristides Dokoumetzidis, Evangelos Karakitsios, Salvatore D’Agate, Alessandro Di Deo, Umberto Villani, Paolo Tieri, Filippo Castiglione, Oscar Della Pasqua

## Abstract

**Background and Objective:** Predicting treatment outcomes in infectious diseases requires accounting for the interplay between drug effects, pathogen dynamics, and host immunity. Integrating pharmacological and immunological approaches into a single simulation environment remains a fundamental challenge in both theory and practice. We aimed to develop and validate a multiscale in silico framework coupling these processes, and to quantify their respective contributions to bacterial clearance.

**Methods:** We present the Drug–Host–Pathogen Interaction (DHPI) framework, combining three independent mechanistic components: a physiologically based pharmacokinetic model of drug disposition, a pharmacokinetic– pharmacodynamic model of drug-induced bacterial killing, and a stochastic agent-based model of the immune response. Continuous concentration profiles are time-averaged onto the agent-based time grid, assigned to bacterial phenotypic states, and converted into per-agent killing probabilities, so that drug-mediated and immune-mediated death events are recorded separately at each step. The framework was applied to simulate symptomatic pulmonary tuberculosis. Phenotype-specific drug-efficacy parameters were inferred using Approximate Bayesian Computation from historical clinical data on eight weeks of 600 mg rifampicin monotherapy, and validated against independent early bactericidal activity data over a disjoint time window.

**Results:** The calibrated framework reproduced the observed decline in bacterial load, and matched reported early bactericidal activity over the first week. In a virtual cohort of symptomatic patients, drug-mediated killing accounted for 81–88% and immune-mediated killing for 12–19% of total bacterial elimination over the 60-day treatment course, while the dormant, granuloma-contained fraction rose from 0.20–0.29 in the first week to 0.85– 0.89 at treatment completion. Over a follow-up of up to 50 years, patients reaching clinical cure had accumulated more memory lymphocytes during treatment than those progressing to clinical failure or death; moreover, the final outcome depended on the immune changes occurring during therapy rather than on the initial disease stage.

**Conclusions:** The results show that the DHPI framework can reproduce treatment dynamics observed in patients and enable the analysis of how therapy reshapes host immune responses and subsequent disease trajectories. By explicitly representing drug–host–pathogen interactions, it provides a mechanistic basis for in silico treatment simulations and for the study of long-term immune consequences of antimicrobial therapy.

## 1. Introduction

Classical pharmacology has traditionally characterised drug efficacy through the direct action of drugs on pathogens. For example, in antibacterial pharmacology, hollow fiber systems, minimum inhibitory concentration assays and time–kill assays quantify killing in the absence of host components [1].

In patients, however, the host immune system (IS) actively modulates bacterial dynamics, influences drug response, and contributes independently to bacterial clearance [2]. Treatment outcome therefore emerges from the interaction of three components—drug, pathogen, and host immunity—rather than from pharmacodynamics alone. This interplay is particularly relevant for host-directed therapies, which aim to modulate immune pathways to enhance microbial clearance or limit immunopathology.

*In vitro* systems, including specialised organ-on-chip platforms [3], are limited in their ability to recapitulate the cellular diversity, spatial organisation, and temporal dynamics of infection in a living host. Despite differences in pathophysiology, *in vivo* animal models offer greater insight into these interactions but are costly, raise ethical concerns, and present translational challenges due to interspecies differences in immune response and pharmacokinetics. Computational biology has developed as a complementary tool, mechanistic models that can integrate knowledge across biological scales, simulate large virtual cohorts, and dissect the contributions of individual system components in a controlled and reproducible setting [4].

Several computational frameworks have addressed parts of this problem. Ordinary differential equation (ODE) models have described immune–pathogen dynamics at the compartmental level [5, 6]. Quantitative systems pharmacology models have incorporated pharmacokinetic–pharmacodynamic (PKPD) representations of antimicrobial activity [7], and agent-based (ABM) or hybrid multiscale models have captured cellular-scale heterogeneity and spatial structure [8–10]. Integration of drug disposition properties and host response has also been explored by coupling a whole-body physiologically based pharmacokinetic (PBPK) model with a population-level immune model calibrated to preclinical PKPD data [11]. Other systems pharmacology models (QSP) have linked immune dynamics of drug exposure in infectious disease settings, although typically without an explicit tissue-resolved PBPK component [12, 13].

Building on these lines of work, our objective is to develop an integrated multiscale framework that combines immune dynamics and drug disposition to quantify the respective contributions of drug-mediated and immunemediated bacterial killing during treatment. In addition to characterising efficacy, the framework further enables the study of how the immune state shaped by therapy determines long-term disease outcome.

It combines three established components: the agent-based immune model C-IMMSIM [14–17], a PBPK model of drug distribution in lung tissue [18], and a PKPD model describing drug-induced bacterial killing [19]. Parameter inference is performed via Approximate Bayesian Computation (ABC) [20, 21].

We applied the proposed framework to tuberculosis (TB), which, among bacterial infections, is characterized by substantial biological and pharmacological complexity. Caused by Mycobacterium tuberculosis (Mtb), TB remains a major global health burden, with 10.7 million new cases reported in 2024 [22]. During infection, the host IS constrains Mtb through coordinated innate and adaptive mechanisms that culminate in the formation of pulmonary granulomas. While these structures limit bacterial dissemination, they also create complex microenvironments that harbour dormant bacilli and restrict drug penetration [23, 24]. Consequently, bacterial clearance during therapy is not a purely pharmacological process but reflects the dynamic competition between immune-mediated and antibiotic action.

To illustrate the framework’s capabilities, we simulate a virtual cohort of adult patients with active pulmonary TB treated with 600 mg rifampicin (RIF) monotherapy, representing the minimal configuration required to isolate drug–immune–pathogen coupling. Although current guidelines recommend multidrug regimens to prevent resistance emergence [22, 25], monotherapy provides a controlled baseline for validating the integrated modelling strategy, with multidrug regimens and host-directed therapeutic scenarios as natural extensions. By quantifying bacterial load, simulations enable the separation of drug-mediated bacterial killing from immune activity. Furthermore, the framework enables the identification of correlations between immune memory trajectories during treatment and long-term disease outcomes, demonstrating its ability to study the coupled dynamics of drug action, host immunity, and bacterial populations.

## 2. Methods

### 2.1. Model architecture

Figure 1 illustrates the structure of the integrated framework and the information exchanged between its components. Each model has been developed and validated independently [14, 15, 19, 26]. The PBPK model provides time-resolved unbound drug concentrations in three lung compartments, which are associated with specific ABM states. Using the PKPD model, these concentrations are converted into bacterial killing rates, to be applied to individual Mtb agents within the ABM at each simulation step.

**Figure 1:**
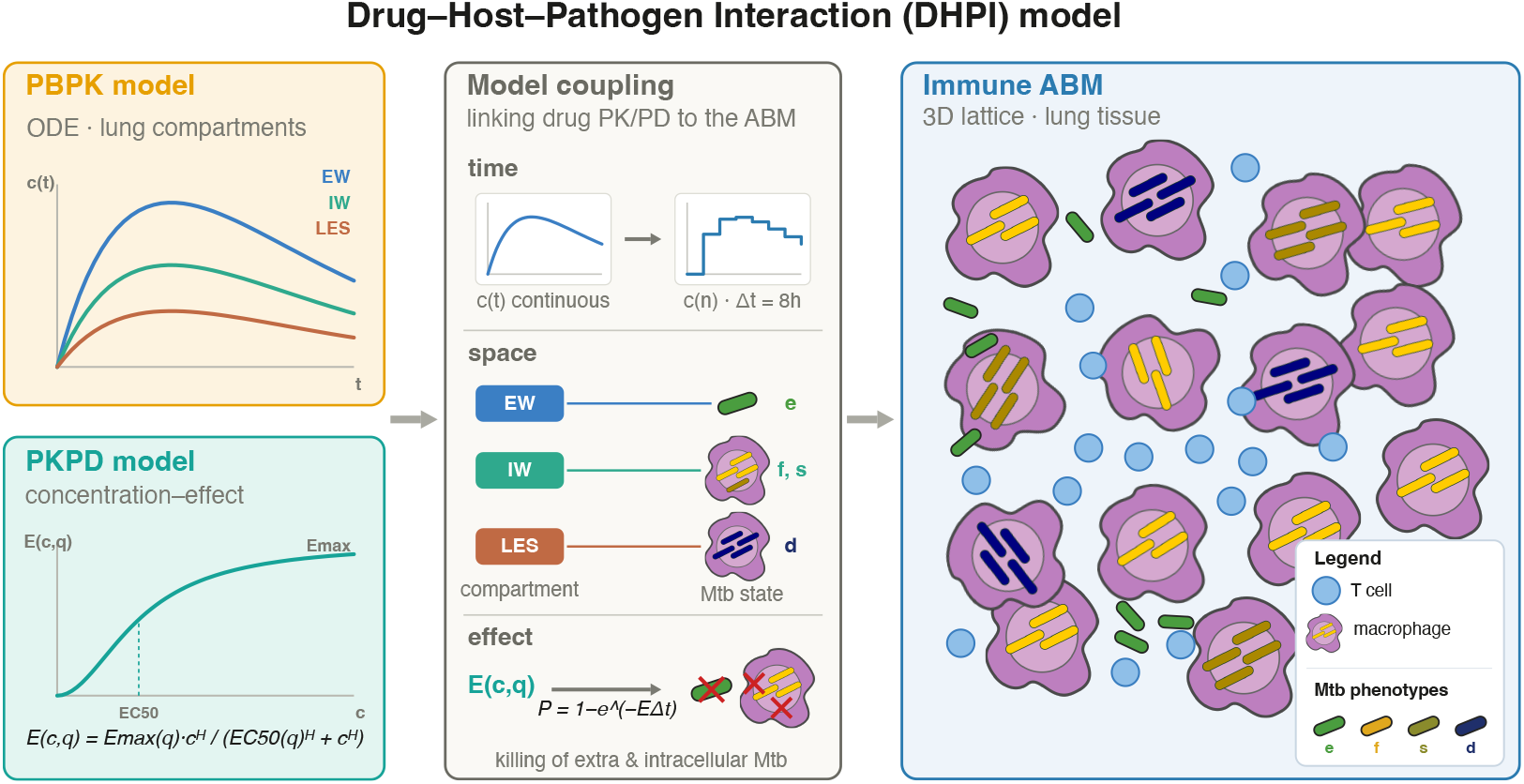
Architecture of the integrated DHPI framework. From left to right, the PBPK model provides continuous-time unbound rifampicin concentrations in the three lung compartments (EW, IW, and LES). The PKPD model converts these concentrations into a drug-induced death rate through the concentration–effect relationship *E*(*c, q*). The resulting outputs are coupled to the C-ImmSim immune agent-based model through three mappings: in time, concentrations are averaged over each 8 h ABM step; in space, each compartment-specific concentration is assigned to the corresponding Mtb phenotype; and in effect, the death rate *E*(*c, q*) is converted into a per-step killing probability *P* (*c, q*) = 1 − exp[−*E*(*c, q*)Δ*t*]. Within the ABM, Mtb agents in four phenotypic states are exposed to both drug-mediated killing and immune-mediated killing.

#### PBPK model of lung drug distribution

Anti-tuberculosis drugs must be transported from the bloodstream to pulmonary lesions and caseous regions where bacteria reside, penetrating not only vascularised but also non-vascularised tissue and diffusing into the necrotic core (caseum). Beyond reaching the lesion, drugs must permeate the Mtb cell envelope to reach intracellular targets at sufficient concentrations and for adequate duration [27]. This distribution to the sites of mycobacterial infection is hindered, which can compromise treatment efficacy. To address this, we employ a PBPK model of RIF distribution in lung tissue [18], validated against in vivo pharmacokinetic data, that integrates drug-specific properties with physiological and anatomical parameters to describe absorption, distribution, metabolism, and elimination in biologically relevant compartments, leveraging micro-dissection homogenate data from distinct lung regions. Using a system of ODEs, the model tracks the time course of unbound RIF concentration across three lung compartments: extracellular water (**EW**), intracellular water (**IW**), and tissue lesion (**LES**), providing for each compartment *k* ∈ {EW, IW, LES} a continuous-time concentration profile 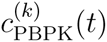.

#### PKPD model of drug-induced bacterial killing

RIF-induced Mtb killing is described through a nonlinear PKPD model characterising antimicrobial activity in an intracellular THP-1 infection system [28]. Bacterial growth and drug-mediated killing were monitored over 10 days under 13 RIF concentration levels ranging from 0.001 to 4 *µ*g/mL (two-fold log_2_ dilutions) using confocal microscopy time-kill assays conducted at Institute Pasteur Lille, France (Eik Hoffmann, Cyril Gaudin; unpublished data). Bacterial dynamics in the absence of drug follow a Verhulst logistic growth model [29] with intrinsic growth rate *k*_gr_ and carrying capacity *B*_MAX_. Drug-mediated killing is incorporated as a first-order process with Hill concentration dependence:

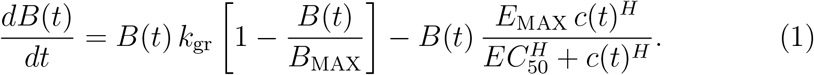

Parameter estimation was performed using the first-order conditional method with interaction [30], as implemented in NONMEM [31]. The estimated parameters *E*_MAX_, *EC*_50_, and *H* represent a population-averaged antibacterial susceptibility, as the in vitro system does not distinguish between intracellular Mtb phenotypes. Phenotype-specific killing rates are resolved through an additional calibration step described in Section 2.3.

#### Immune ABM (C-IMMSIM)

The immune component is based on C-IMMSIM [14–17, 32], extended to model TB infection dynamics [26]. The ABM represents a portion of lung tissue as a three-dimensional lattice populated by immune entities and Mtb agents, evolving in discrete time with step Δ*t* = 8 h, which is adequate to describe the agent interactions represented in the ABM. Immune agents include macrophages, dendritic cells, natural killer cells, B lymphocytes (including plasma cells), CD4^+^ T helper cells, CD8^+^ cytotoxic T cells, antibodies, and cytokines. Interactions follow stochastic, rule-based mechanisms governing recruitment, activation, cytotoxicity, phagocytosis, antigen presentation, and memory formation.

Within the ABM, Mtb is represented as an agent assuming four phenotypic states [32]: extracellular replicating (*e*), fast-growing intracellular (*f* ), slow-growing intracellular (*s*), and dormant/non-replicating (*d*) (Figure 2).

**Figure 2:**
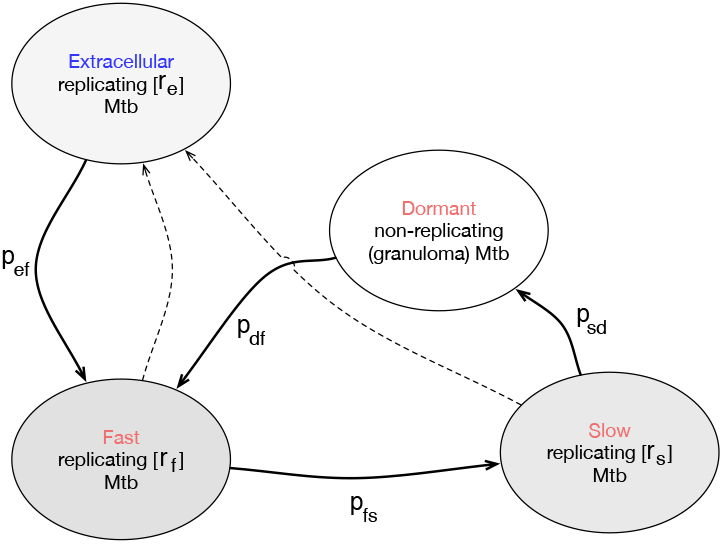
Stochastic finite state machine which represents the state diagram of each Mtb agent. The arrows indicate the possible state transitions, each associated with a calibrated probability [26].

At the onset of infection, non-phagocytosed bacteria are in the extracellular state, characterised by low motility and low replication rate. Upon phagocytosis, bacteria enter the fast-growing intracellular state, where replication increases, reflecting the favourable intracellular environment of the macrophage when it fails to eliminate the pathogen. From there, bacteria may transition to a slow-growing phenotype characterised by reduced metabolism and prolonged intracellular survival. Extracellular, fast- and slow-growing bacteria are collectively referred to as replicating Mtb. Replicating intracellular bacteria may exit host cells upon macrophage lysis, while a subset instead enters a dormant state, in which replication ceases and no immune interaction takes place—a representation of granuloma containment without sterilisation [23, 33]. Dormant bacteria may later reactivate, reproducing the phenomenology of granuloma rupture [34, 35], which may lead to active infection.

#### Integration of PBPK, PKPD and ABM

The PBPK model provides continuous-time concentration profiles, whereas the ABM evolves on a discrete time grid with step Δ*t* = 8 h. To provide the ABM with a representative drug concentration at each simulation step, PBPK concentrations are mapped to the ABM grid by time-averaging over each interval:

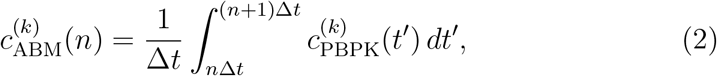

where *k* ∈ {EW, IW, LES} and *n* denotes the ABM time-step index.

The three PBPK compartments are mapped to the spatial organisation of the ABM and the Mtb phenotypic states as follows (Figure 3). The **EW** concentration is assigned to extracellular replicating bacteria (*q* = *e*). The **IW** concentration is assigned to fast- and slow-growing bacteria (*q* ∈ {*f, s*}) residing within non-infected macrophages. Finally, the **LES** concentration is assigned to fast- and slow-growing bacteria (*q* ∈ {*f, s*}) harboured by infected macrophages, defined as those carrying more than 10 intracellular bacteria [26]. Note that while **IW** and **LES** target the same phenotypic states, they differ in the drug concentration assigned, reflecting the reduced RIF penetration into lesioned tissue. As a first approximation, dormant bacteria (*q* = *d*) are not sensitive to drug effect.

**Figure 3:**
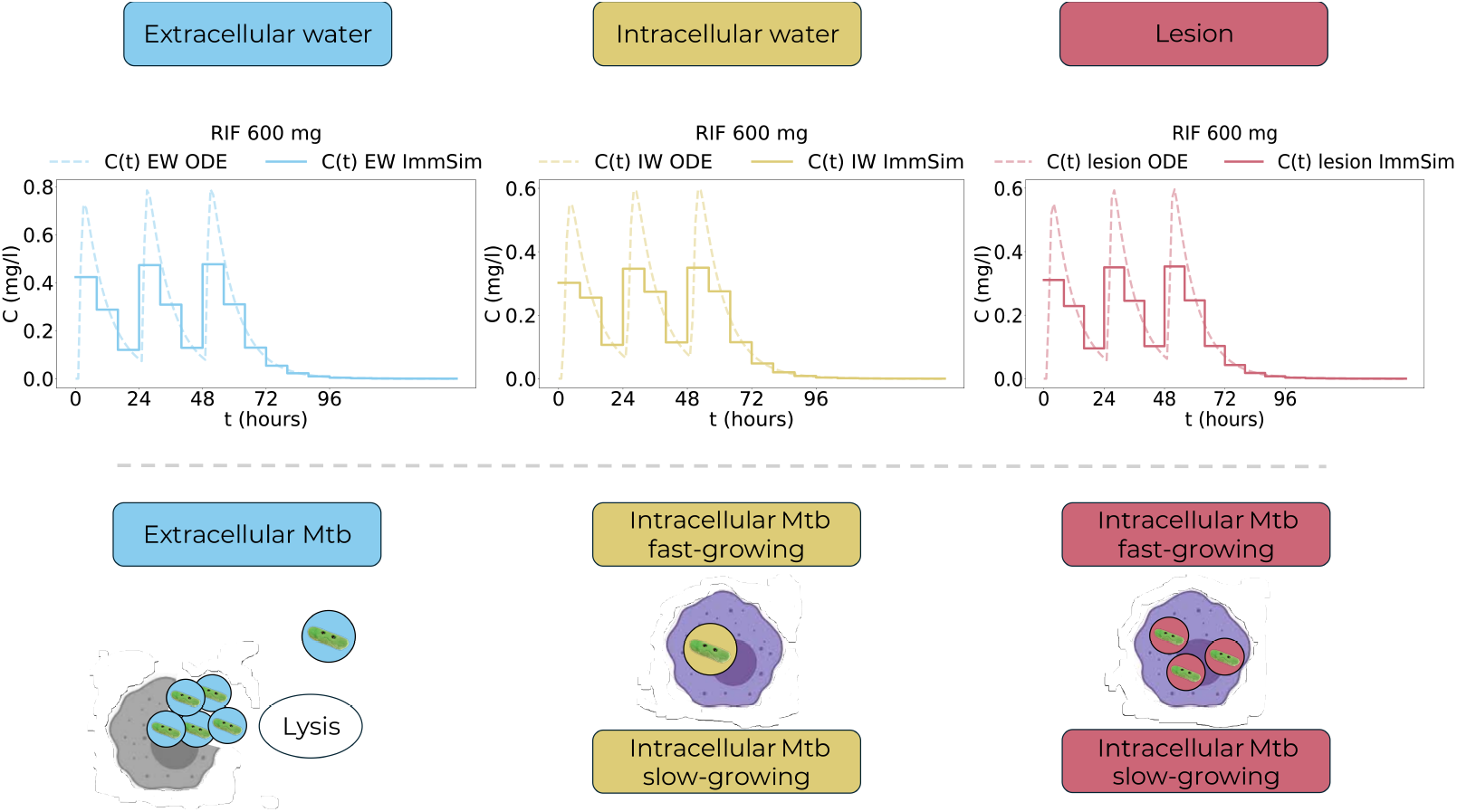
Mapping of PBPK compartments and drug concentrations into the ABM. Top panels: time course of unbound RIF concentration in the extracellular water (EW), intracellular water (IW), and lesion (LES) compartments. The dashed lines exemplify the continuous PBPK ODE solutions for 3 dosing events, while the solid stepwise lines correspond to the time-averaged concentrations mapped onto the discrete ABM time grid (Δ*t* = 8 h). Bottom panels: Mtb phenotypic states in the ABM and their spatial association with PBPK compartments. Colours indicate the compartment–state correspondence used for drug exposure mapping.

For integration within the ABM, only the concentration–effect component of the PKPD model is retained, as bacterial replication and immune-mediated clearance are already represented mechanistically at the agent level. The population-averaged parameters *E*_MAX_, *EC*_50_, and *H* estimated in vitro are extended to a phenotype-specific relationship: for a bacterium in state *q* exposed to local unbound concentration *c*, the drug-induced death rate is:

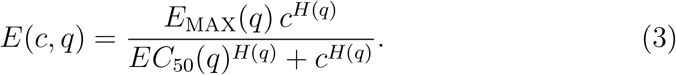

Here *E*(*c, q*) represents a population-level instantaneous death rate. To implement drug effect at the level of individual agents within the ABM, this rate is converted into a per-agent, per-step elimination probability via the standard survival function:

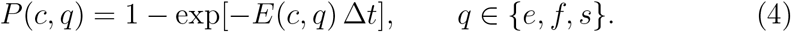

At each simulation step, every replicating bacterium in state *q* is independently removed with probability *P* (*c, q*) according to the concentration assigned through the mapping above. The resulting concentration–effect relationship *E*(*c, q*) and the corresponding killing probability *P* (*c, q*) for extracellular and intracellular Mtb under 600 mg RIF are shown in Appendix B. A key feature of this coupling is that immune-mediated and drug-mediated bacterial death events are recorded separately at each ABM step, enabling quantification of the respective contributions of the IS and pharmacological treatment to bacterial killing throughout the treatment period and beyond.

### 2.2. Study Workflow

This study is organised in two conceptually distinct phases (Figure 4). Phase I is devoted to the generation of a virtual cohort and to the estimation of the therapy-related parameters of the multiscale framework. In this phase, simulations of the ABM are used to generate a heterogeneous untreated population of virtual patients and to identify the subset of symptomatic disease trajectories. These trajectories are defined as the time evolution of the host–pathogen system (bacterial load, immune response, and clinical state). A subset of symptomatic trajectories is then used to calibrate the phenotype-specific drug-effect parameters of the integrated framework by comparison with clinical data on RIF monotherapy.

**Figure 4:**
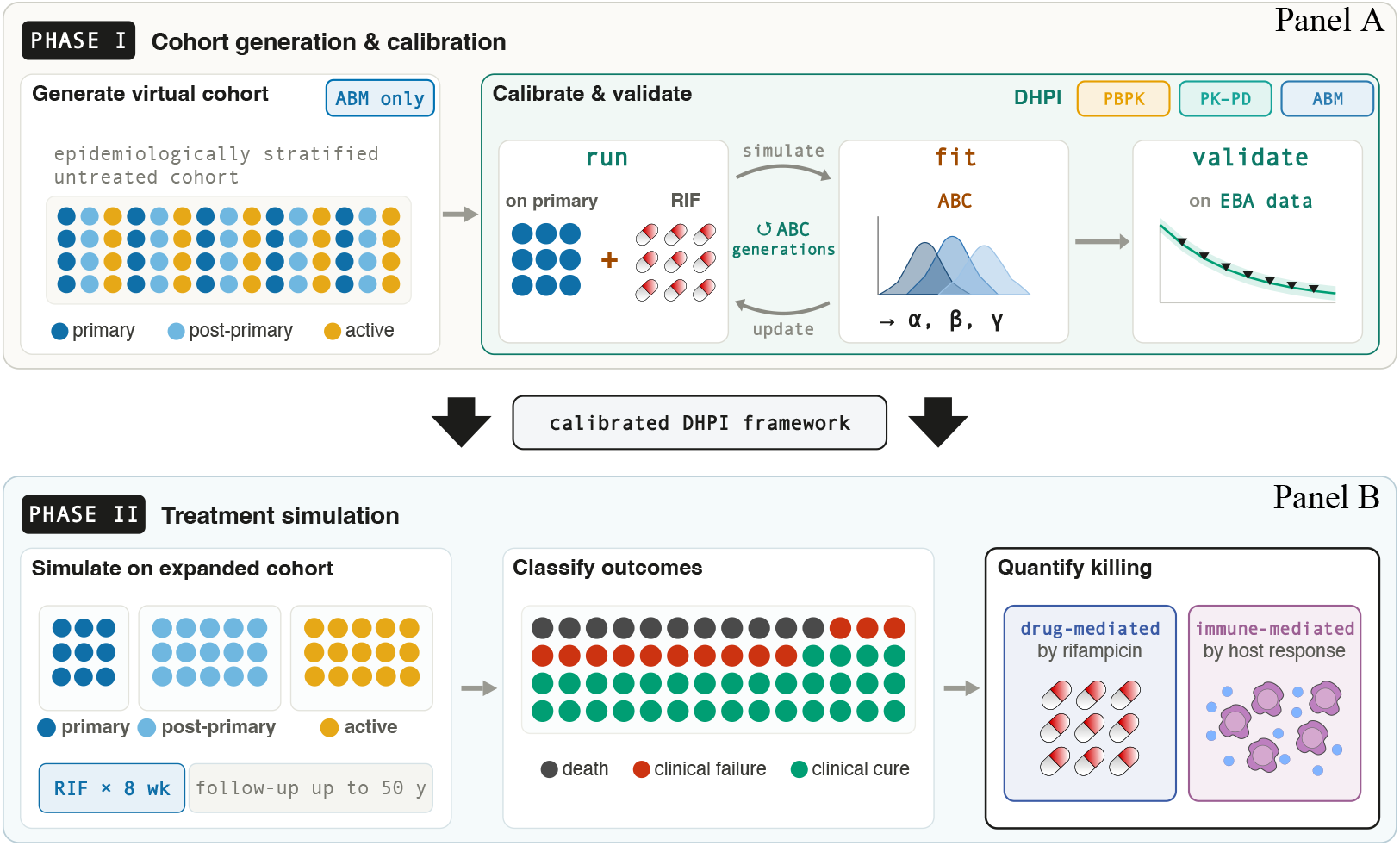
Two-phase in silico experimental workflow. **Panel A (Phase I):** ABM is used to generate the virtual cohort, patients are classified by disease outcome. Symptomatic patients are selected and used to calibrate therapy-related parameters. **Panel B (Phase II):** The DHPI framework is used to simulate treatment dynamics for an expanded virtual cohort. The relative contribution of immune and drug-mediated bacterial killing is quantified.

In Phase II, once the framework parameters are calibrated, the model is applied to a larger cohort of symptomatic patients to simulate treatment dynamics, analyse long-term disease trajectories, and quantify the respective contributions of drug-mediated and immune-mediated bacterial killing.

#### Phase I – Cohort generation and framework calibration

*Virtual cohort generation.* A virtual cohort of heterogeneous individuals is generated instantiating ABMs characterised by a different parametrization (Section 2.1). Each virtual patient represents an immunologically healthy adult defined by a distinct initial immune configuration, including variability in immune-cell homeostasis, repertoire diversity, MHC composition. Virtual patients are simulated for up to 50 years after infection.

ABM parameters are calibrated so that the distribution of outcomes aligns with the quantitative targets of the epidemiological tree (Figure A.8), which summarises reported population-level frequencies of TB outcomes including clearance, latency, active disease, relapse, and death. Cohort generation and epidemiological calibration are inherited from Mastrostefano et al. [26] and described in more detail in Appendix A.

Disease progression of a virtual individual is categorised on the basis of thresholds on the replicating bacterial load *L_rep_*: symptomatic if it exceeds THR_active_ = 1×10^5^ bacteria at time *t_s_*, and deceased if it exceeds THR_death_ = 7 ×10^7^. Symptomatic patients are stratified as *primary TB* (fatal within two years without therapy), *post-primary TB* (fatal after two years), or *active TB* (symptomatic but non-fatal).

#### Framework calibration

Primary TB patients (*N* = 61, as identified in Mastrostefano et al. [26]) constitute the subset used to estimate the therapy-related parameters of the integrated framework. These patients are therefore re-simulated under the full multiscale model, with treatment initiated at *t_s_*. Phenotype-specific drug-efficacy parameters governing drug-induced bacterial killing in the ABM are estimated via ABC by matching simulated bacterial-load dynamics during eight weeks of 600 mg RIF monotherapy against clinical data [36]. The resulting parameter estimates are validated on an independent clinical dataset on a disjoint temporal window [37]. These simulations are used exclusively for calibration and validation of the framework parameters.

#### Phase II – Treatment simulations and mechanistic analysis

Once the frame-work parameters are calibrated, the model is applied to an expanded symptomatic cohort composed of 61 primary, 100 post-primary, and 100 active TB virtual patients. Each patient receives 600 mg RIF monotherapy once daily for eight weeks starting at *t_s_*, and simulations are followed for up to 50 years.

These simulations are used to analyse treatment trajectories, quantify the relative contributions of drug-mediated and immune-mediated bacterial killing, and investigate the long-term immune response associated with different treatment outcomes. Post-treatment trajectories are classified as *death* (*L*_rep_ *>* THR_death_ after treatment initiation), *clinical failure* (symptomatic recurrence without reaching THR_death_), *clinical cure* (no recurrence but persistent bacterial load), or *recovered* (full bacterial clearance). No virtual patient achieved the recovered outcome, consistent with the persistence of dormant bacteria not targeted by rifampicin.

### 2.3. Model calibration and validation

#### Calibration parameters

The experimental data used to estimate the PKPD model do not allow for the characterisation of PKPD relationships in different intracellular bacterial subpopulations; the parameter estimates *E*_MAX_, *EC*_50_, and *H* therefore represent a population-averaged antibacterial susceptibility over a mixed intracellular population that may consist of both fast- and slow-growing bacteria in unknown and time-varying proportions. Within the ABM, however, the concentration–effect relationship must be specified separately for each phenotypic state *q* ∈ {*e, f, s*}. To this end, we model the aggregate in vitro efficacy *E*(*c, i*) against intracellular bacteria as:

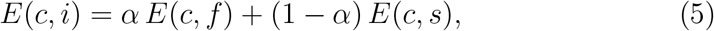

where *E*(*c, f* ) and *E*(*c, s*) are the efficacies on fast- and slow-growing bacteria, respectively, and *α* ∈ [0, 1]. This formulation approximates the drug effect on different Mtb phenotypes by averaging its impact over the duration of the in vitro assay, as capturing the full intracellular dynamics during the experiments is beyond the scope of this work.

Because the experiments report only a population-averaged antibacterial effect on a heterogeneous intracellular bacterial population, the contributions of fast- and slow-growing bacteria cannot be identified separately from the data. We therefore introduce a structural assumption to decompose the aggregate efficacy into phenotype-specific effects. As a first approximation, we assume that the efficacies on the two phenotypes are proportional to each other, introducing the parameter *β* ∈ R^+^:

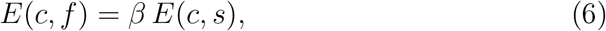

where *β <* 1 indicates greater efficacy on slow-growing bacteria and *β >* 1 indicates the opposite. Finally, we introduce the constant *γ >* 0, common to all bacterial phenotypes, which represents a global scale factor accounting for systematic differences between the controlled in vitro environment and the physiological context represented in the ABM, for example, the different drug exposure times in vitro, with parameter estimates based on 24-hour experiments, compared to the 8-hour temporal resolution of the ABM.

Combining Equations (5), (6), and (3), the complete set of state-specific drug efficacies applied within the ABM is:

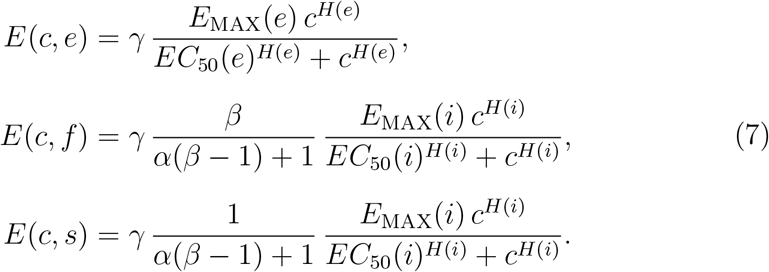

Note that while the spatial compartment determines which PBPK concentration profile *c*(*t*) is assigned to a given bacterium, it does not alter the functional form of the efficacy: at fixed concentration, *E*(*c, q*) depends only on the phenotypic state *q* ∈ {*e, f, s*}.

#### Calibration data

Clinical data on RIF monotherapy are limited in the recent literature. Available studies typically report early bactericidal activity (EBA) over only the first one to two weeks of treatment [37]. On the other hand, real-world data on longer monotherapy durations are generally only available from the period prior to the adoption of the standard multi-drug protocol [22]. Monotherapy is therefore considered here as a controlled experimental setting that allows the pharmacological effect of a single drug to be isolated for model calibration.

For the calibration of (*α, β, γ*) we use the dataset reported by Gyselen [36], consisting of mean weekly Gaffky scores measured on sputum smear microscopy in *n* = 10 patients over eight weeks of 600 mg RIF monotherapy. The Gaffky score is a semi-quantitative index of bacillary density that can be converted into an estimated bacterial count per millilitre of sputum using published mappings [38].

Since only mean Gaffky scores are reported, additional processing is required to reconstruct an experimental target trajectory; the full procedure is described in Appendix C.

The output of this procedure is an individual-level normalised bacterial load,

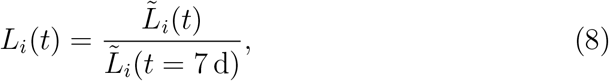

where *L̃_i_*(*t*) denotes the estimated active (replicating, non-dormant) bacterial burden of individual *i*, and the denominator normalises to the value one week after treatment initiation. Following [39, 40], sputum smear counts are assumed to reflect the extracellular plus intracellular (excluding dormant) Mtb population and to be directly proportional to the active bacterial load. Under this approximation, ⟨*L*^(^*^E^*^)^(*t*)⟩ and ⟨*L*^(^*^M^*^)^(*t*)⟩ are directly comparable, where ⟨·⟩ denotes averaging over experimental data for *L*^(^*^E^*^)^ and over model simulations for *L*^(^*^M^*^)^. The resulting values of ⟨*L*^(^*^E^*^)^(*t*)⟩ are reported in Table C.2 (Appendix C).

We acknowledge that these assumptions are simplifying. Sputum smear measurements provide only an indirect and semi-quantitative estimate of the underlying bacterial burden, and the proportionality assumption is introduced to enable calibration. More precise experimental measurements explicitly discriminating between bacterial subpopulations would allow this mapping to be refined in future work.

#### Calibration procedure

The framework parameters (*α, β, γ*) are inferred using the ABC method [20], implemented with pyABC [21]. ABC is adopted because the stochastic, multi-scale nature of the framework precludes explicit evaluation of a tractable likelihood function. Parameter inference is therefore performed by iteratively sampling from prior distributions and retaining parameter sets that produce simulated trajectories sufficiently close to the experimental data according to a predefined distance metric.

Prior to the full ABC run, a preliminary exploratory analysis over a coarse 45-point parameter grid was performed to identify values yielding clinically realistic treatment trajectories, thereby excluding regions of parameter space associated with unrealistic rapid sterilisation or negligible drug effect (Appendix D).

At each ABC iteration, a sampled triplet (*α, β, γ*) is evaluated by running *N* = 61 independent ABM simulations.

Model–data distance is quantified using the symmetric mean absolute percentage error (SMAPE):

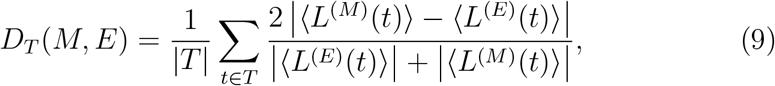

where *T* = {7, 14, 21, 28, 35, 42, 49, 56} (days) is a set of time points and |*T* | denotes its cardinality. SMAPE is chosen because the normalised bacterial load spans multiple orders of magnitude during treatment and approaches zero at later time points; unlike standard percentage error metrics, SMAPE remains well-defined in this regime and bounds the distance in [0, 2]. Alternative distance metrics, including mean absolute percentage error and root mean square deviation, were evaluated in a preliminary analysis (Appendix D) and yielded consistent results.

The accepted sample size per generation is fixed at *S* = 500. At each generation, the tolerance threshold *ɛ* is updated to the 0.4 quantile of the distances obtained in the previous generation, following the default adaptive scheme implemented in pyABC. The calibration is run for 8 generations. Convergence is assessed by visual inspection of the stability of the marginal posterior distributions and of the evolution of *ɛ* across generations (Figure 5). ABC yields an approximate joint posterior distribution for (*α, β, γ*). Point estimates are taken as posterior means, and the full posterior is propagated to quantify parameter uncertainty.

**Figure 5:**
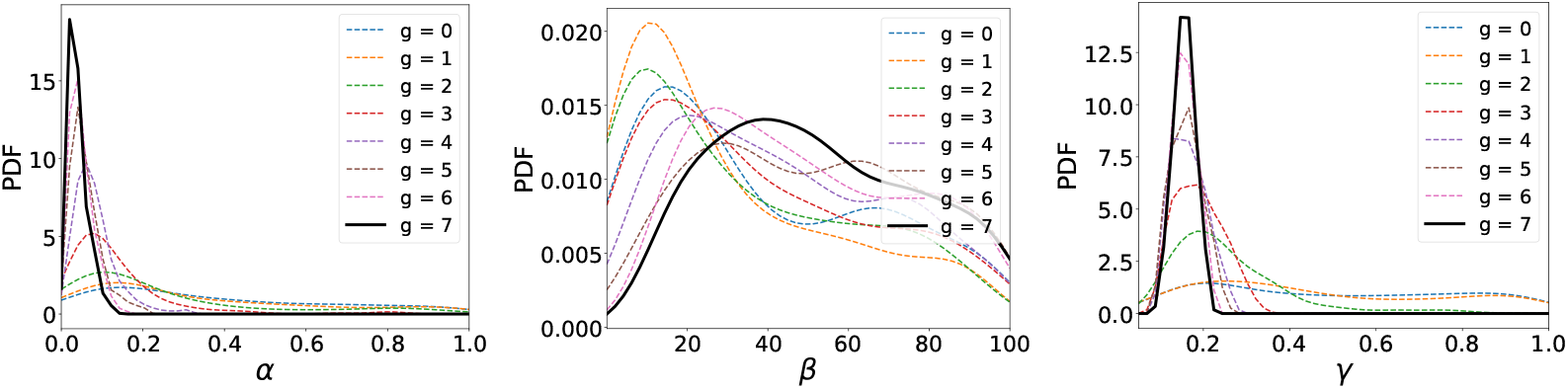
Evolution of the marginal posterior distributions of *α*, *β*, and *γ* across ABC generations *g* = 0*, . . . ,* 7. Each panel shows the progressive concentration of the posterior toward the final estimate. The solid black line corresponds to generation *g* = 7, representing the converged result of the calibration. Left: fraction of fast-growing bacteria in the in vitro population (*α*). Centre: ratio of drug efficacy on fast-versus slow-growing bacteria (*β*, log scale). Right: global in vitro-to-ABM scaling factor (*γ*).

## 3. Results

The DHPI framework provides a mechanistic setting in which drug exposure, bacterial phenotypes, and host immune responses are represented within a single simulation environment. This allows treatment dynamics to be analysed in terms of the coupled action of pharmacological and immunemediated processes. The results presented below first assess the validation of the integrated model against clinical data, and then use the framework to quantify the relative contributions of drug and IS to bacterial killing and to analyse the resulting long-term disease outcome, including how therapy-induced changes in the immune response influence post-treatment disease trajectories.

### Validation

We first assess whether the integration of three independently developed models can be calibrated in a principled and reproducible manner from clinical data, and whether the calibrated system reproduces observations outside the calibration window (i.e. extrapolation).

In the TB case study, ABC inference yields joint posterior distributions for the phenotype-specific drug-efficacy parameters (*α, β, γ*) (Figure 5). Calibration is performed on eight weeks of 600 mg RIF monotherapy data, using the SMAPE distance defined in Eq. 9. After convergence of the ABC procedure, the posterior means are ⟨*α*⟩ = 0.04 ± 0.02, ⟨*β*⟩ = 52 ± 25, and ⟨*γ*⟩ = 0.16 ± 0.02. These values correspond to the mean of the joint posterior distribution, while the full posterior samples are retained for uncertainty propagation in downstream simulations.

The posterior of *α* is centered near, but not at, zero. The inferred values are not negligible in terms of model output, as small variations in this parameter induce substantial changes in the predicted probability at simulated concentrations (e.g., approximately a twofold reduction in death probability as *α* increases from 0 to 0.04).

The posterior of *β* is centered well above unity, indicating a clear separation between the inferred efficacies assigned to fast- and slow-growing phenotypes within the phenotype-resolved extension of Eq. 7. The posterior distribution of *γ* remains confined to a narrow interval around its mean value and does not approach the boundaries of the prior range. In multi-component computational frameworks, the absence of extreme rescaling factors indicates that the coupled model components operate on mutually compatible scales, without requiring compensatory parameter inflation to reconcile differences in temporal resolution or mechanistic granularity.

Validation is performed on an independent temporal window not used for calibration, namely the first week of therapy (days 0–7), with EBA data reported by Brake et al. [37], which provide daily decline rates of −0.111 (mean), −0.194 (lower 95% CI), and −0.028 (upper 95% CI) log_10_ CFU mL*^−^*^1^ day*^−^*^1^. From these rates, three reference trajectories are reconstructed over days 0– 7 and compared against the mean simulated active bacterial load ⟨*L*^(^*^M^*^)^(*t*)⟩ of the virtual patients with active infection, under the calibrated parameter values (Figure 6). Agreement between reconstructed EBA-based trajectories and simulated dynamics indicates that the integrated framework generalises beyond the calibration interval while preserving internal model consistency.

**Figure 6:**
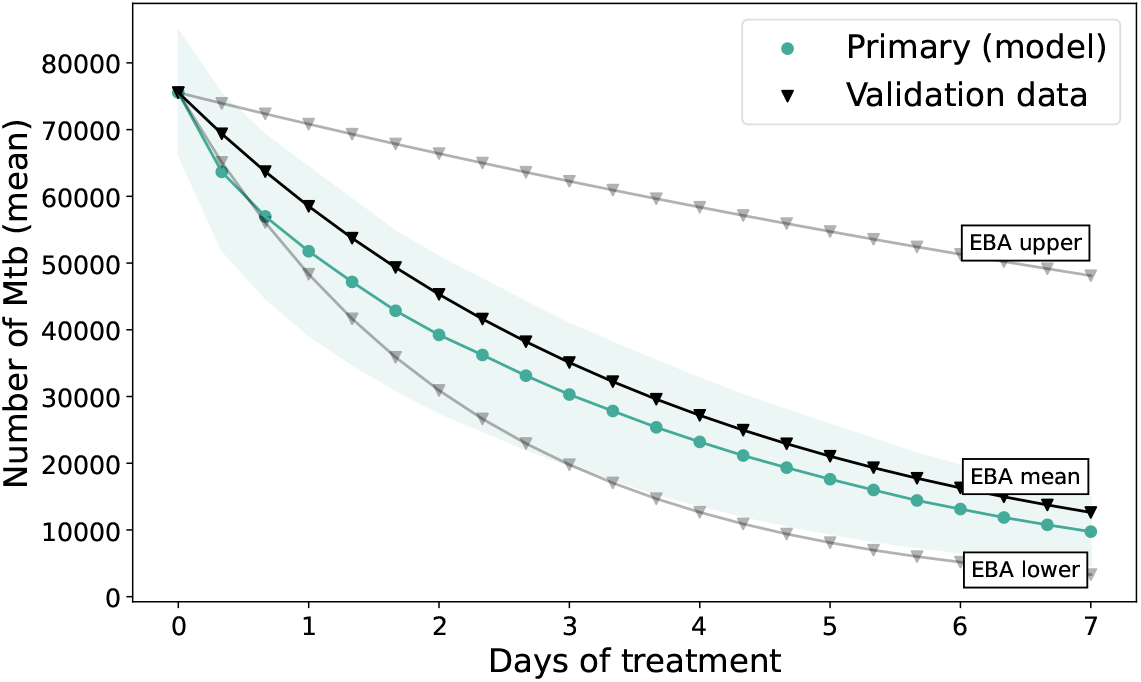
Validation of the calibrated parameters against independent short-term data. Green circles: mean active bacterial load of the 61 primary virtual patients during the first week of 600 mg RIF monotherapy simulated under the calibrated parameter values (shaded area: standard deviation). Triangles: bacterial load trajectories reconstructed from the EBA values reported in Brake et al. [37]. Black triangles: mean EBA; grey triangles: lower and upper 95% CI bounds.

### IS contribution to clearance

At each ABM time step, replicating bacteria are removed either through the drug-dependent killing probability (Eq. 4) or through immune-mediated events. Total elimination therefore satisfies *R*_tot_(*t*) = *R*_IS_(*t*) + *R*_RIF_(*t*), where *R*_IS_(*t*) and *R*_RIF_(*t*) denote the number of bacteria killed in [*t* − Δ*t, t*] by the IS and by RIF, respectively. This parametrisation makes it possible to define the relative immune contribution:

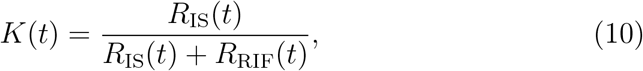

which quantifies the fraction of net bacterial reduction attributable to host-mediated killing. Importantly, *K*(*t*) is a direct simulation output. In the tuberculosis case study, this decomposition indicates that immune-mediated killing accounts for 9–16% of total elimination during the first treatment week and 12–19% over the full 60-day course, with the complementary fraction attributable to RIF (Table E.3). These values are consistent across outcome stratification, showing that the relative pharmacological contribution during treatment is not the primary source of long-term divergence within the simulated cohort. Because tuberculosis is characterised by granuloma-mediated containment and dormancy, an additional TB-specific descriptor is introduced to quantify the surviving bacterial population. The dormant fraction:

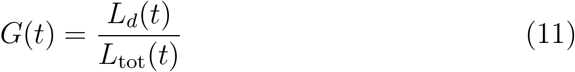

measures the proportion of bacteria contained within granulomas. Unlike *K*(*t*), which captures elimination, *G*(*t*) captures containment dynamics. During therapy, *G*(*t*) increases from approximately 0.20–0.29 in the first week to 0.85–0.89 by the end of treatment (Table E.3), indicating clearance of fast and slow growing bacteria and a progressive shift of the population toward dormancy.

### Long-term dynamics after treatment

A distinct feature of the framework concerns the relationship between treatment dynamics and long-term outcome. Because treatment does not reset the ABM state, the immune configuration reached at therapy completion serves as the initial condition for long-term dynamics. Post-treatment dynamics are followed for up to 50 years and classified as death, clinical failure, or clinical cure (Section 2.2).

To analyse immune behaviour at the level of individual reactivation episodes, we consider granuloma rupture events occurring at times *t_i_* and compute two event-level descriptors:

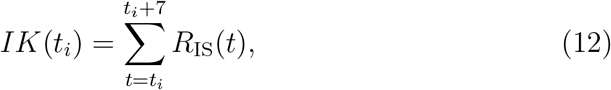

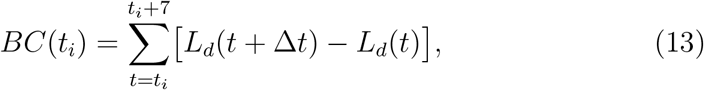

representing, respectively, the total immune-mediated killing and the net change in granuloma-contained bacteria over a one-week period following rupture, chosen as a biologically relevant timescale for immune activation [41]. Each rupture event therefore defines a point in the (*BC, IK*) plane.

Figure 7 compares the joint distribution of (*BC*(*t_i_*)*, IK*(*t_i_*)) before and after treatment.

**Figure 7:**
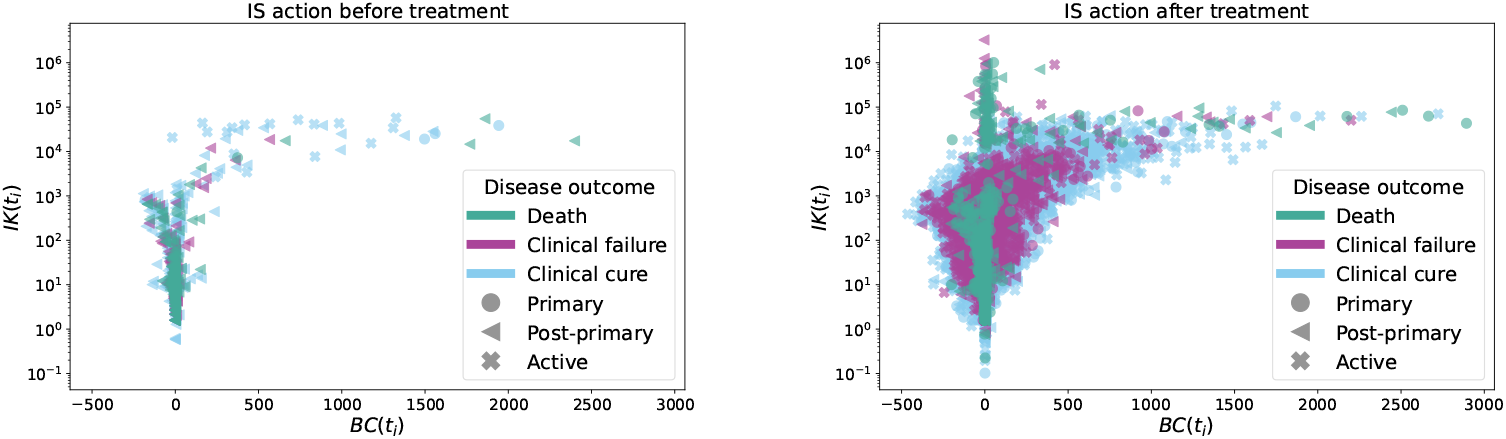
Event-level immune response to granuloma rupture. On the *x*-axis: net granuloma formation *BC*(*t_i_*) (Eq. 13). On the *y*-axis: immune-mediated killing *IK*(*t_i_*) (Eq. 12). Each point corresponds to a granuloma rupture event. Colours indicate post-treatment outcome (death, clinical failure, clinical cure). Marker shapes denote pre-therapy stratification (primary, post-primary, active). Left panel: rupture events before treatment. Right panel: rupture events after treatment.

Before treatment, the distributions of bacterial containment and immune-mediated killing overlap across all trajectory classes, with no pattern linking immune response to eventual outcome. After treatment, a clear separation emerges. Fatal trajectories are characterised by failure to re-establish bacterial containment (*BC* ≈ 0), whereas cured trajectories remain confined to bounded regions of the (*BC, IK*) plane (Figure 7). This separation is not predicted by pre-therapy classification into primary, post-primary, or active disease, but emerges as a consequence of treatment itself. The mechanistic basis for this divergence lies in differential accumulation of memory lymphocytes during therapy administration. Cured patients develop substantially larger increases in memory B, CD4^+^, and CD8^+^ T cell counts over the course of treatment, with statistically significant differences relative to both death and clinical failure groups (Table 1). This suggests that immune remodelling occurring during therapy administration, rather than pre-treatment disease classification, is the primary determinant of whether the host can mount an effective response to subsequent reactivation events.

**Table 1:**
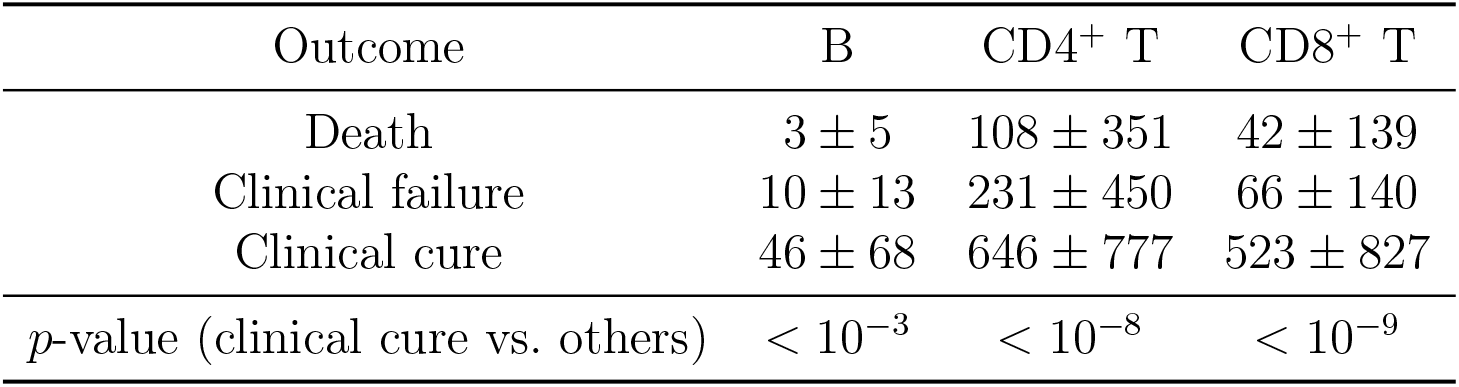
Differences in memory immune cell counts between the last and first day of pharmacological treatment, averaged for each outcome after treatment (death, clinical failure, clinical cure). The last row reports the largest *p*-value from Welch’s *t*-test comparing the clinical cure outcome mean with the means of the other outcomes.

| Outcome | B | CD4<sup>+</sup> T | CD8<sup>+</sup> T |
| --- | --- | --- | --- |
| Death | $3 \pm 5$ | $108 \pm 351$ | $42 \pm 139$ |
| Clinical failure | $10 \pm 13$ | $231 \pm 450$ | $66 \pm 140$ |
| Clinical cure | $46 \pm 68$ | $646 \pm 777$ | $523 \pm 827$ |
| $p$-value (clinical cure vs. others) | $< 10^{-3}$ | $< 10^{-8}$ | $< 10^{-9}$ |

## 4. Conclusion

We have presented a multiscale, data-informed in silico framework integrating three independently validated mechanistic components: a PBPK model of drug distribution in lung tissue, a PKPD model of drug-induced bacterial killing, and a stochastic agent-based model of the immune response, within a unified simulation environment. The integrated system is calibrated via Approximate Bayesian Computation and validated on independent clinical data. Our results demonstrate that mechanistically heterogeneous models can be jointly parameterised in a principled and reproducible manner from real-world observations.

In the tuberculosis case study, applied to 600 mg rifampicin monotherapy, the framework quantifies that approximately 81–88% of bacterial elimination during treatment is drug-mediated and 12–19% is immune-mediated. Because drug- and immune-driven killing events are explicitly tracked at each simulation step, this separation arises from disentanglement of the mechanisms rather than from post-hoc inference. The same structural continuity allows the immune configuration reached at therapy completion to determine long-term dynamics, revealing that treatment modifies the immune system behaviour subsequent granuloma rupture events and that memory lymphocyte accumulation during the dosing window correlates with long-term outcome, a prognostic signal accessible during therapy itself.

Current limitations include the restriction to monotherapy and the absence of adherence variability, resistance emergence, and comorbid conditions, as well as the use of time-averaged drug effects and proportional efficacy assumptions to characterize Mtb phenotypes. Extension to multidrug regimens and larger clinical datasets will allow further assessment of the robustness and generalisability of the framework. The modular architecture provides a structured basis for mechanistically informed in silico treatment simulations in infectious diseases where drug distribution, pathogen dynamics, and host immunity interact across scales.

## Acknowledgments

This work received support from the Innovative Medicines Initiative 2 Joint Undertaking (grant No. 853989; http://www.imi.europa.eu).

This work reflects only the author’s views, and the JU is not responsible for any use that may be made of the information it contains.

## CRediT authorship contribution statement

**Alessandro Ravoni:** Data curation, Formal analysis, Investigation, Methodology, Software, Visualization, Writing – original draft, Writing – review & editing. **Enrico Mastrostefano:** Conceptualization, Formal analysis, Investigation, Methodology, Software, Supervision, Writing – original draft, Writing – review & editing. **Davide Moretti:** Data curation, Methodology, Software, Visualization, Writing – review & editing. **Elia Onofri:** Data curation, Methodology, Software, Writing – review & editing. **Francesca Pelusi:** Methodology, Software, Visualization, Writing – review & editing. **Aristides Dokoumetzidis:** Data curation, Methodology, Supervision. **Evangelos Karakitsios:** Data curation, Formal analysis, Methodology. **Salvatore D’Agate:** Data curation, Formal analysis, Methodology. **Alessandro Di Deo:** Data curation, Formal analysis, Methodology, Software. **Umberto Villani:** Data curation, Formal analysis, Methodology, Software. **Paolo Tieri:** Funding acquisition, Investigation, Supervision, Writing – review & editing. **Filippo Castiglione:** Funding acquisition, Investigation, Software, Supervision, Writing – review & editing. **Oscar Della Pasqua:** Conceptualization, Investigation, Supervision, Writing – review & editing.

## Ethics Statement

No ethical approval was required for this study.

## Appendix A. The C-ImmSim Agent Based Model

C-ImmSim is a stochastic agent-based model of the innate and adaptive immune response, implemented as a system of cellular automata evolving on a three-dimensional lattice [14, 15]. Details about the model and examples of its applications can be found in previous publications [16, 17, 32]. This section summarises the aspects of the platform that are relevant to the present work.

### Entities, lattice and time

The lattice maps the volume of tissue in which the immune response takes place; in the present work, a portion of lung, where pulmonary infection is established.

Simulated entities are of two kinds. *Cells* are autonomous agents that retain their individual experience throughout their simulated lifespan. The innate ones are macrophages, dendritic cells and natural killer cells; the adaptive ones are B lymphocytes, antibody-producing plasma cells, CD4^+^ T helper and CD8^+^ cytotoxic T lymphocytes. *Molecules* are simpler entities characterised by their number, position and binding site, and include antigens, antibodies, immune complexes and cytokines.

Cells move at random across the lattice and interact only with entities occupying the same site, so that all interactions are local. One time step corresponds to Δ*t* = 8 h, of the order of a lymphocyte mitotic cycle. Cell numbers are kept at homeostasis, and each simulation starts from an immunologically naive host.

### Molecular specificity as binary strings

C-ImmSim is polyclonal: it does not track a single representative clone but an entire repertoire of competing ones. Binding sites are encoded as binary strings of fixed length. B- and T-cell receptors, MHC molecules and antigen epitopes are therefore all represented in the same formalism, and the potential repertoire size grows exponentially with the string length.

Binding is governed by complementarity: a 0 in one string faces a 1 in the other. The number of complementary positions—equivalently, the Hamming distance between the two strings—measures the strength of a match, and affinity grows monotonically with it, above a threshold below which no binding occurs. Recognition is therefore graded rather than all-or-none.

The binary strings are an abstraction: they capture specificity combinatorially, and not through the actual amino-acid sequences of peptides, receptors and MHC alleles. This is sufficient for the repertoire-level quantities used in this work, such as lymphocyte counts, but not for questions about which individual epitopes are recognised. An extension of the platform recovers the sequences by coupling the simulator to molecular binding prediction tools [16].

### Lymphocyte generation and repertoire selection

Lymphocytes are generated in a bone marrow compartment, where each cell draws its receptor at random. B lymphocytes enter the circulation directly. T lymphocytes first transit through a thymus compartment, where a set of strings defines the self of the simulated individual and any cell reacting against it is removed. Each virtual patient therefore has its own randomly drawn, self-tolerant repertoire, and this is one of the sources of the immune heterogeneity of the virtual cohort.

### Innate and adaptive dynamics

Antigen-presenting cells, namely B cells, macrophages and dendritic cells, take up and digest antigen, and display the resulting peptides on MHC molecules. Peptides on class I molecules activate CD8^+^ cytotoxic cells, those on class II activate CD4^+^ helper cells. B lymphocytes that engage antigen and receive help from activated T helper cells proliferate clonally and may mutate their receptor at division, so that clones of progressively higher affinity emerge over successive rounds of selection.

Immunological memory is modelled as a cell state rather than as a separate differentiation branch. A cell acquires it by taking part in successful interactions: each of them extends the half-life of the cell, so that lymphocytes repeatedly engaged by antigen persist while the others decay. The memory compartment therefore grows with the magnitude and the duration of the response.

### Representation of Mtb and phenotypic states

Mtb is modelled as an agent assuming different bacterial phenotypes, with transitions between states occurring as probabilistic events once per simulation time step (Figure 2).

As outlined in Section 2.1, at the beginning of infection, Mtb exists in an extracellular state, i.e. non-phagocytosed, characterised by low motility and a low replication rate. Once engulfed by a macrophage, the bacteria transition into a fast-growing intracellular phenotype, in which the replication rate increases. This reflects the ability of Mtb to exploit the intracellular environment when the host cell fails to effectively eliminate the pathogen.

From this stage, bacteria may further differentiate into a slow-growing intracellular phenotype, characterised by reduced metabolic activity and prolonged intracellular persistence. In this state, replication occurs at a lower rate, which is thought to contribute to immune evasion and long-term survival within the host.

Eventually, bacteria replicating within macrophages can exit upon host-cell lysis, while some enter a dormant state in which they do not replicate and are not detected by the IS. The dormant state captures key aspects of granulomas, i.e., containment of bacteria without sterilisation. Dormant bacteria may later reactivate, potentially leading to disease relapse and reproducing the phenomenology of granuloma rupture [34, 35].

Macrophages containing more than 10 intracellular bacteria are classified as *infected* and interpreted as representing lesion-like conditions [26]. This threshold introduces a spatial compartmentalisation in the ABM corresponding to the stratification of drug exposure: the extracellular region (targeting extracellular Mtb), the intracellular region (targeting fast- and slow-growing Mtb within non-infected macrophages), and the lesioned region (targeting fast- and slow-growing Mtb within infected macrophages). Bacteria in the dormant state are not affected by drug treatment.

### Epidemiological classification and calibration targets

The epidemiology of TB has been extensively characterised through clinical and population studies. A spectrum of disease states has emerged from this body of work, shaped by varying bacterial loads and immunological activity, in particular questioning the binary distinction between active TB and latent TB. This spectrum is often shaped by granulomas, complex structures composed of immune cells and tissue, including regions of lesion formation and necrosis, within which Mtb is in a dormant, non-replicating state that allows the immune system (IS) to contain the infection.

In this context, Mastrostefano et al. [26] proposed a classification of TB history that emphasises a spectrum of probabilities rather than discrete categories of TB outcomes. The resulting epidemiological tree (Figure A.8) summarises the quantitative relationships, derived from the literature, among clinical states. Each branch corresponds to a distinct clinical or subclinical pathway passing through various possible states characterised by different bacterial loads and immunological activity: clearance, latency, active disease, relapse, and death. The tree serves both as a conceptual synthesis and as a quantitative reference for calibration: it defines the target frequencies that untreated virtual patients generated by the ABM must reproduce.

**Figure A.8:**
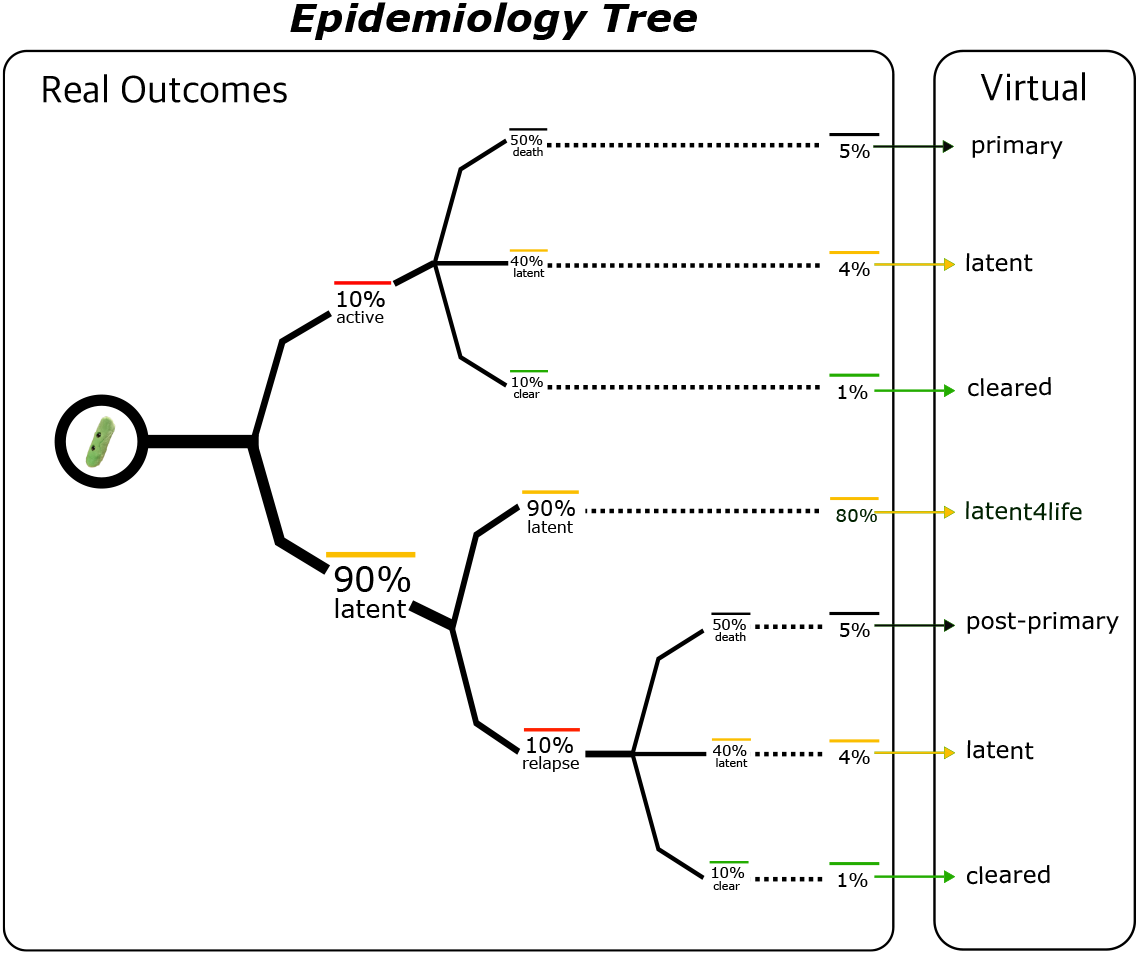
Left panel: frequency of outcomes of TB represented as a tree (distribution of outcomes). Each branch is associated with a frequency, expressed as a percentage. The leaves are the final outcomes with the associated risk. Right panel: simulated disease states associated with each outcome.

The calibration process uses a disease state classification based on threshold values for the load of replicating bacteria. A virtual patient is considered symptomatic (active TB) when the replicating load exceeds THR_active_ = 1 × 10^5^ bacteria, and deceased if it exceeds THR_death_ = 7 × 10^7^ bacteria. Patients who develop symptoms are further divided into three groups reflecting the ability of the IS to contain the infection in the absence of therapy: *primary TB* patients experience fatal outcomes within the first two years after infection; *post-primary TB* patients experience fatal outcomes after the first two years; *active TB* patients develop symptomatic but non-fatal disease.

### Immune heterogeneity and virtual cohort generation

Each virtual patient represents an immunologically healthy adult characterised by a distinct initial immune configuration, including variability in immune-cell homeostasis, the diversity of the immune repertoire, the composition of MHC complexes, and the stochastic events that unfold during the simulation. This variability ensures that the virtual cohort spans a realistic range of immune competence with respect to Mtb infection. Untreated simulations are carried out for up to 50 years after infection. Model parameters are calibrated so that the resulting distribution of simulated outcomes aligns with the quantitative targets defined by the epidemiological tree.

It is important to note that the model does not consider direct interactions between drug effects and the IS: the only implemented action of RIF is the reduction of Mtb counts. Although recent studies suggest an antagonistic relationship between antitubercular drugs and the IS, this interplay remains poorly understood. We reserve the possibility of investigating this aspect further in future work.

## Appendix B. Concentration–effect and killing probability

Figure B.9 shows the concentration–effect function *E*(*c, q*) (left panel) and the corresponding discrete-time killing probability *P* (*c, q*) (right panel) for extracellular and intracellular Mtb. The curves are obtained using the PKPD parameters *E*_max_, *EC*_50_, and *H*estimated from in vitro time-kill experiments (unpublished data, Institut Pasteur Lille).

## Appendix C. Smear data and estimation of *L*(*t*)

In Gyselen [36] the author reports the Gaffky score measured on the smear of 10 patients treated for 8 weeks with 600 mg RIF monotherapy. We use Figure 1 shown in Gyselen [36] to estimate the average Gaffky score measured as a function of time, which we report in the Table C.2. The Gaffky score is a discrete score ranging from 1 (lowest Mtb load) to 10 (highest Mtb load), each value providing an estimation of the number of bacteria in smear; we also note that several studies suggest the observed population comprises different phenotypes of both extracellular and intracellular bacteria [39, 40]. In Table 1 of Okada [38], the author reports estimates of the number of bacteria measured in the sputum as a function of the assigned Gaffky score across different experiments, although these estimates are affected by a large degree of uncertainty. In particular, by averaging the values measured in the various experiments reported in Okada [38], we obtain the following estimates for the number of bacteria in smears corresponding to Gaffky scores of 1, 2, 3 and 4:

**Figure B.9:**
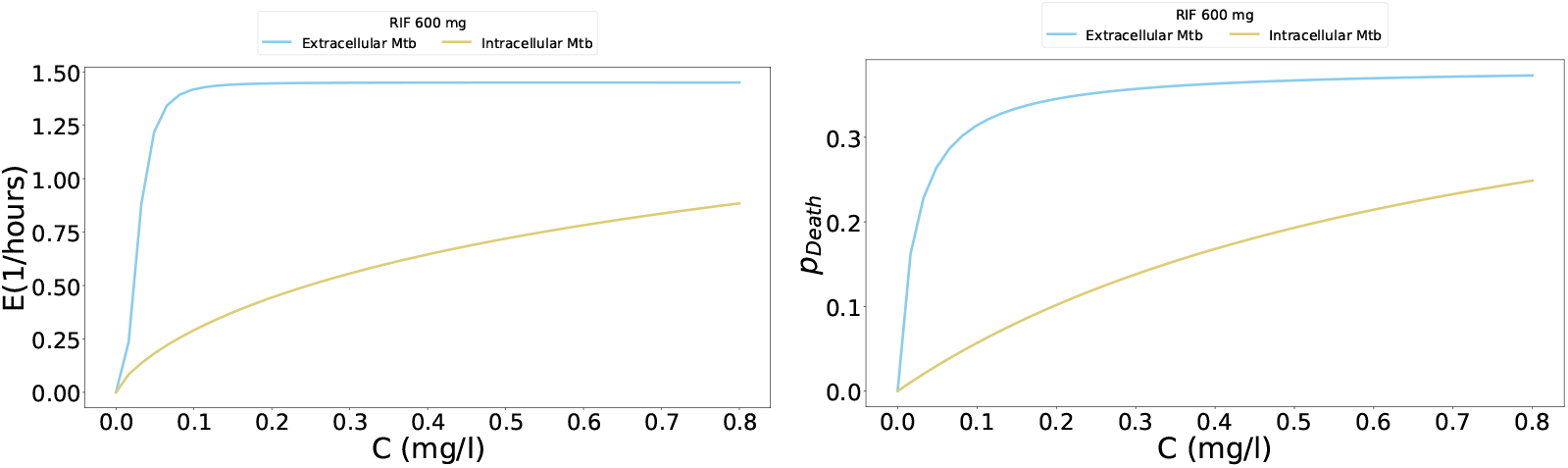
RIF 600 mg effect in the ABM. Left panel: estimated drug efficacy (Equation 3) for extracellular and intracellular Mtb. Right panel: corresponding killing probability (Equation 4). Parameter values for *EC*_50_, *E_MAX_* , and *H* for RIF used in this study are estimated from in vitro time-kill experiments (unpublished data, Institut Pasteur Lille).

- Gaffky score 1 → (2.3 ± 1.8)10^3^ bacteria/mL;
- Gaffky score 2 → (3.3 ± 4.3)10^5^ bacteria/mL;
- Gaffky score 3 → (1.7 ± 1.1)10^6^ bacteria/mL;
- Gaffky score 4 → (2.4 ± 1.7)10^6^ bacteria/mL;

We show in Figure C.10 the resulting estimated Mtb load measured in Gyselen [36]. Since the values reported in Gyselen [36] are averaged over multiple patients and thus represent continuous quantities, data processing is needed to combine the average Gaffky scores with the bacterial counts associated with each discrete score. We identify different approaches to address the problem: the first one is to define a function that maps each continuous value within the relevant Gaffky score range to the corresponding bacterial count, with its parameters estimated by fitting the results reported in Okada [38]. However, a preliminary analysis showed that such a function must be non-linear, which leads to a significant amplification of the already large experimental data uncertainty.

Therefore, we decided to adopt a different strategy, specifically by reconstructing a population of individuals with associated Gaffky score measurements at each time point of interest, from which the average number of bacteria can be estimated. To do so, we assume that each experimental Gaffky score *µ_t_* obtained at time *t* corresponds to the mean of a normal distribution N(*µ_t_, σ*). We further assume that it is *σ* = 0.25, such that the Gaffky scores of the population described by N(*µ_t_, σ*) fall within the interval *µ_t_* ±1 with a probability of ≈ 0.95. We thus reconstruct the Gaffky score values at different time points over a population of 10 individuals by sampling from N(*µ_t_, σ*) and rounding the values to the nearest integer, subsequently discarding any negative values. The sampled values can be mapped to bacterial counts in smear based on the average values obtained from Okada [38], with the additional assumption that a Gaffky score of zero corresponds to the absence of bacteria.

This approach allow us to collect *L̃_i_*(*t*) for a population of *i* = 1*, . . .* 10 individuals, under the assumption that the extracellular plus intracellular (excluding dormant) Mtb load in TB patients is directly proportional to the bacterial counts in smear.

Finally, we introduce the normalized quantity *L_i_*(*t*) = *L̃_i_*(*t*)*/L̃_i_*(*t* = 7), where *L*(*t* = 7) is the number of bacteria measured one week after the start of treatment, and compute the average ⟨*L*(*t*)⟩ over the population. Under the proportionality constraint, ⟨*L*(*t*)⟩ is representative of the mean values of both the experimental data on the number of bacteria measured in the smear, the bacterial load in human patients and the simulated bacterial load. The values found of ⟨*L*^(^*^E^*^)^(*t*)⟩ obtained from experimental data are reported in Table C.2.

## Appendix D. Exploratory analysis

We perform a preliminary exploratory analysis to investigate the behavior of the integrated model for extreme values of the parameters controlling the effect of the therapy. Specifically, we consider 45 parameter triplets obtained by combining the following values:

- *α* ∈ [0, 0.5, 1], corresponding to assuming that the experimental efficacy for intracellular bacteria was measured on a population consisting solely of slow-growing Mtb, solely of fast-growing Mtb, and an equal mix of the two phenotypes, respectively;
- *β* ∈ [0.01, 0.1, 1, 10, 100], corresponding to assuming a drug action ranging from being 100 times more effective on slow-growing Mtb than on fast-growing ones, to 100 times more effective on fast-growing Mtb than on slow-growing ones;
- *γ* ∈ [0.01, 0.1, 1], corresponding to assuming a model granularity for bacterial cells ranging from a factor of 100 to full correspondence between a simulated agent and a real cell.

**Table C.2:** Time course of Mtb in patients treated with 600 mg RIF monotherapy. First column: days since the start of treatment with 600 mg RIF. Second column: average Gaffky score values measured in 10 patients with active TB. The values are estimated from Figure 1 shown Gyselen [36]. The error in the values corresponds to the estimated observational error. Third column: corresponding values of the function ⟨*L*^(^*^E^*^)^(*t*)⟩, which describes the average number of Mtb over time in humans, normalized to the value at one week after treatment. The reported errors correspond to the propagation of experimental error in the estimation of the bacterial count in the smear associated with each Gaffky score.

| Days since treatment start | Gaffky score | $\langle L^{(E)}(t) \rangle$ |
| --- | --- | --- |
| 7 | $2.8 \pm 0.02$ | $1.0 \pm 0.7$ |
| 14 | $1.25 \pm 0.02$ | $(0.4 \pm 1.3) \times 10^{-1}$ |
| 21 | $0.76 \pm 0.02$ | $(1.1 \pm 1.2) \times 10^{-3}$ |
| 28 | $0.75 \pm 0.02$ | $(1.2 \pm 1.2) \times 10^{-3}$ |
| 35 | $0.62 \pm 0.02$ | $(0.5 \pm 1.2) \times 10^{-3}$ |
| 42 | $0.68 \pm 0.02$ | $(1.1 \pm 1.2) \times 10^{-3}$ |
| 49 | $0.43 \pm 0.02$ | $(0.1 \pm 1.1) \times 10^{-3}$ |
| 56 | $0.56 \pm 0.02$ | $(0.7 \pm 1.2) \times 10^{-3}$ |

**Figure C.10:**
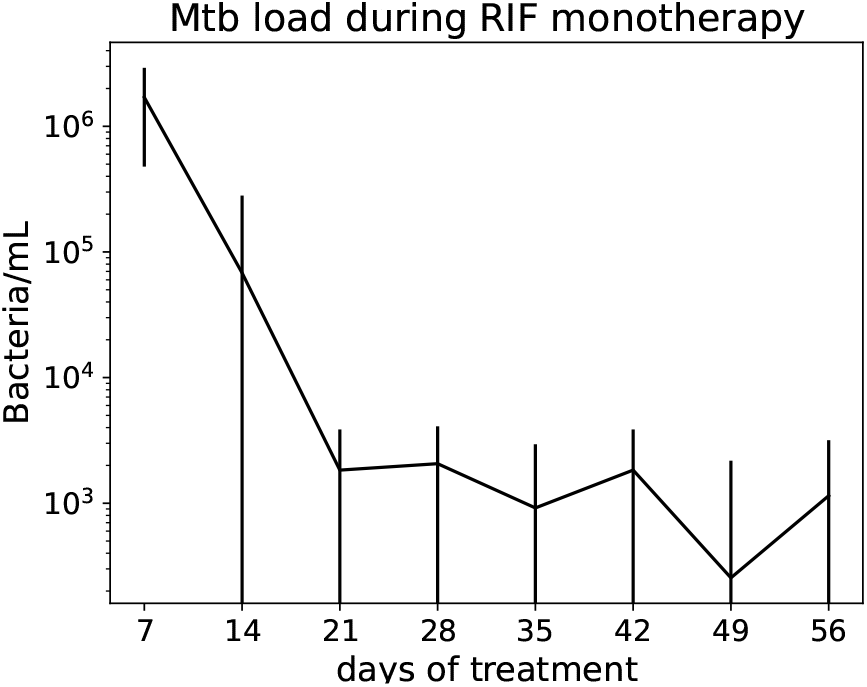
Plot of the estimated Mtb load measured in smear of patients treated with RIF monotherapy. On the x axis: time from the start of treatment (days). On the y axis: Mtb load averaged over 10 patients. The data shown are obtained by combining the mean bacterial load measured in the smear of patients treated with RIF monotherapy in terms of Gaffky score (from Gyselen [36]) with the estimates of the bacterial load associated with the various Gaffky scores (from Okada [38]), as described in Section Appendix C

For each triplet (*α, β, γ*) we apply the protocol described in Section 2.2 to a population of *N* = 61 virtual patients who develop symptoms within the first year after infection onset. The deviation *D*(*T* ) from experimental results was quantified primarily using SMAPE; however, alternative metrics, such as mean absolute percentage error and root mean square deviation, produced consistent results (data not shown).

The outcomes are shown in the Figure D.11. In particular, we observe that the minimum distance of *D*(*T* ) ≈ 1.047 is achieved for *α* = 0, *β* = 100, *γ* = 0.1 and that, in general, the distance tends to be higher for values of *γ* = 0.01. We also note that no clear trend can be identified for the tested values of *α* and *β*, indicating that their range should not be restricted in the final calibration phase.

Moreover, we highlight that, for *γ* ∈ [0.01, 0.1], the cumulative amount of *_t∈T_ L̃_M_* (*t*) at the experimental measurement time points is consistently higher in the simulations compared to the available data. This suggests that the simulated combined effect of therapy and immune response is less effective than what is observed in real patients.

Conversely, in few triplets with *γ* = 1, the opposite occurs. In this case, we observe that therapy leads to a complete absence of extracellular bacteria before the end of the first week after treatment and throughout the entire treatment period in a significant proportion of the *N* virtual patients. We refer to this outcome as quick clearance. This result suggests that the simulated therapy is perhaps overly effective and therefore unrealistic.

We also observe that, for *γ* ∈ [0.1, 1], a substantial fraction of patients exhibits complete elimination of extracellular bacteria, followed by their reappearance later during treatment, which results in a non-zero number of extracellular Mtb at the end of therapy. We refer to this outcome as clinical failure. In our model, clinical failure is triggered by the transition of Mtb from the dormant state to the fast-growing state. This clinically relevant situation should occur, but not affect too many treated patients. These findings suggest that the range of *γ* for ABC calibration should be restricted to values close to [0.1, 1] (we actually perform the calibration by setting *γ* in the range [0.05, 1]).

The percentages of quick cleared (Q) and clinical failure (R) patients associated with each triplet are shown in Figure D.11. Note that, by definition, the two outcomes are mutually exclusive.

**Figure D.11:**
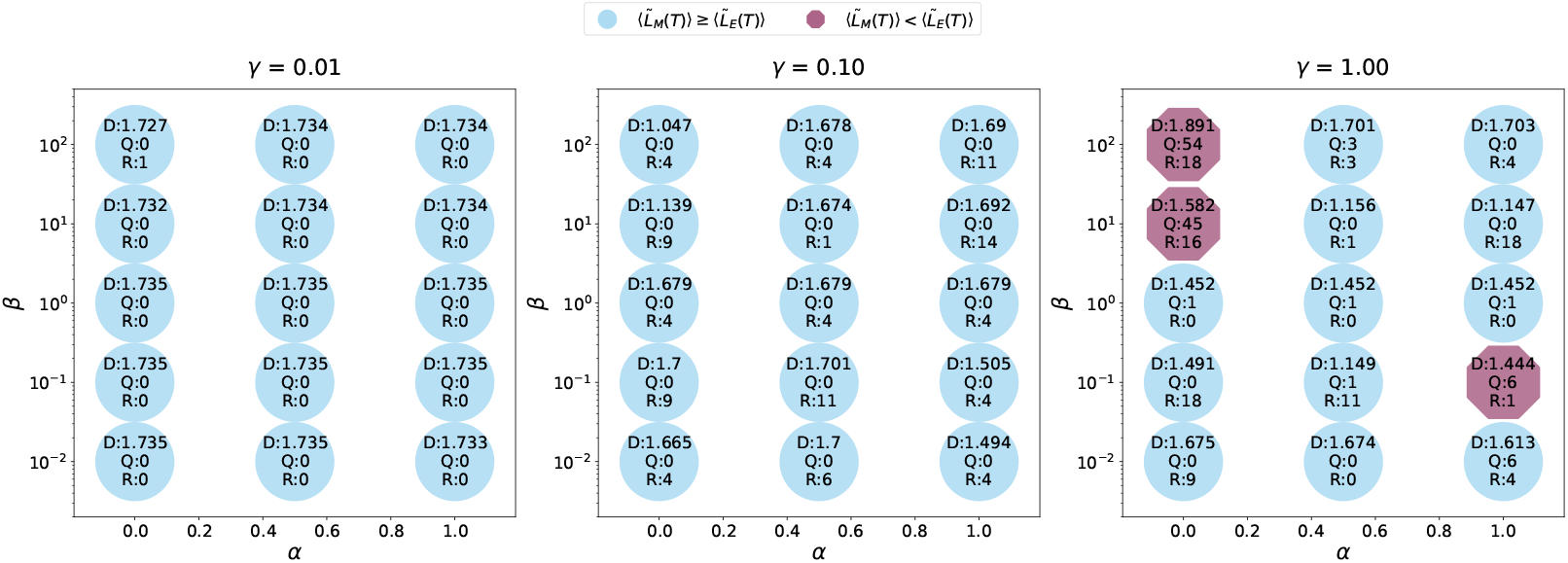
Scatter plot of distance *D*(*T* ) obtained from the exploratory analysis. From left to right: increasing *γ* values. On the x axis: *α*. On the y axis: *β*. Text labels on each point indicate: (D) the distance *D*(*T* ), computed as the difference between the experimental ⟨*L_E_*(*t*)⟩ and the reproduced ⟨*L_M_* (*t*)⟩ averaged across *N* = 61 virtual patients, according to Equation 9; (Q) the number of quick cleared simulations, as defined in Section Appendix D; (R) the number of simulations with clinical failure, as defined in Section Appendix D. Cyan circles: Σ*_t∈T_* ⟨*L_M_* ⟩ −⟨*L_E_*⟩ ≥ 0. Purple octagons: Σ*_t∈T_* ⟨*L_M_* ⟩ −⟨*L_E_*⟩ *<* 0.

## Appendix E. Outcome of treatment

In this section we report a table and some figures of results obtained by administering RIF 600 mg monotherapy to symptomatic virtual patients. Table E.3 shows the mean values of *K*(*t*) and *G*(*t*) across the subgroups of virtual patients, for both the first week and the entire duration of treatment. Figure E.12 shows the number of dormant Mtb and the bacteria killed by the IS and by the drug during the entire treatment in symptomatic virtual patients.

**Figure E.12:**
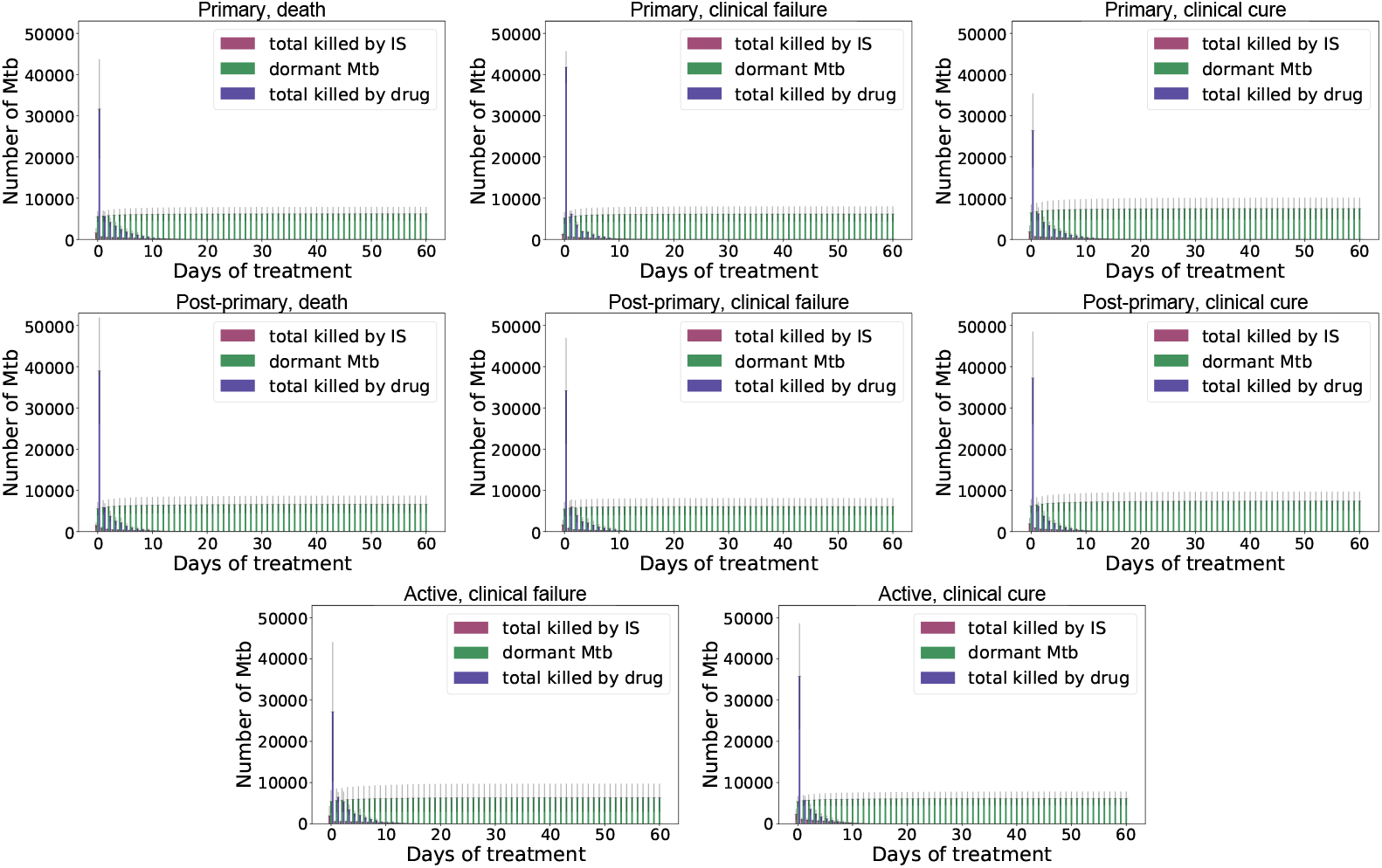
Quantification of IS and drug contributions during the entire treatment of symptomatic (primary, post-primary, active) patients. First row: primary patients. Second row: post-primary patients. Third row: active patients. Left column: death outcome (not present for active patients). Centre column: clinical failure. Right column: clinical cure. On the *x*-axis: days of treatment. On the *y*-axis: number of Mtb killed by IS (purple bars), number of Mtb killed by RIF 600 mg (blue bars), number of dormant Mtb (green bars). Each bar shows the mean value calculated over virtual patients within each subgroup; grey lines show the corresponding standard deviations.

**Table E.3:** Values of quantities *K*(*t*) and *G*(*t*) as defined in Section 3, computed for the first week and for the entire duration of treatment. The values shown are the mean ± standard deviation calculated across each subgroup in which the virtual patients have been classified.

| Disease outcome | $K(7 \text{ days})$ | $K(60 \text{ days})$ | $G(7 \text{ days})$ | $G(60 \text{ days})$ |
| --- | --- | --- | --- | --- |
| death (primary) | $0.11 \pm 0.02$ | $0.13 \pm 0.03$ | $0.22 \pm 0.07$ | $0.86 \pm 0.02$ |
| death (post-primary) | $0.10 \pm 0.02$ | $0.14 \pm 0.02$ | $0.28 \pm 0.09$ | $0.87 \pm 0.02$ |
| clinical failure (primary) | $0.09 \pm 0.02$ | $0.12 \pm 0.02$ | $0.28 \pm 0.06$ | $0.88 \pm 0.02$ |
| clinical failure (post-primary) | $0.10 \pm 0.02$ | $0.13 \pm 0.03$ | $0.26 \pm 0.11$ | $0.87 \pm 0.03$ |
| clinical failure (active) | $0.10 \pm 0.04$ | $0.15 \pm 0.04$ | $0.20 \pm 0.08$ | $0.85 \pm 0.03$ |
| clinical cure (primary) | $0.11 \pm 0.02$ | $0.15 \pm 0.03$ | $0.22 \pm 0.07$ | $0.86 \pm 0.02$ |
| clinical cure (post-primary) | $0.12 \pm 0.02$ | $0.16 \pm 0.03$ | $0.28 \pm 0.08$ | $0.88 \pm 0.02$ |
| clinical cure (active) | $0.16 \pm 0.05$ | $0.19 \pm 0.04$ | $0.29 \pm 0.12$ | $0.89 \pm 0.02$ |

